# Inflammation-enhanced synapse-specific phagocytosis by adult APP microglia in a microfluidic neuron–microglia co-culture model

**DOI:** 10.1101/2025.08.19.671013

**Authors:** Dolores Siedlecki-Wullich, Anne-Marie Ayral, Lukas Iohan, Célia Lemeu, Valérie Buiche, Karine Blary, Julien Chapuis, Fanny Eysert, Delphine Beury, Myriam Delacre, David Hot, Takahiro Masuda, Klaus-Peter Knobeloch, Marco Prinz, Jean-Charles Lambert, Devrim Kilinc

## Abstract

Microglia play a critical role in synapse remodeling and neuroinflammation, both of which are dysregulated in Alzheimer’s disease (AD). However, most in vitro models rely on neonatal or immortalized microglia, limiting their relevance to adult pathophysiological context. Here, we present a compartmentalized microfluidic co-culture platform that enables spatially controlled interactions between primary cortical neurons and adult microglia from wild-type (WT) and APP-transgenic mice. This system allows precise functional analysis of microglia–synapse interactions under defined inflammatory conditions. Upon lipopolysaccharide (LPS) stimulation, APP microglia exhibited exaggerated morphological activation, elevated IL-1β secretion, and selectively increased engulfment of synaptic material. In contrast, phagocytosis of non-specific substrates such as pHrodo™ Zymosan remained unchanged, suggesting a substrate-specific enhancement of microglial phagocytic activity. Blocking the complement receptor CD11b abolished the LPS-induced increase in synaptic uptake, confirming the role of complement-dependent pathways. Transcriptomic profiling revealed robust inflammatory responses in both genotypes, with selectively heightened expression of proinflammatory genes in APP microglia, consistent with a primed immune phenotype. Importantly, increased synaptic uptake occurred without measurable loss of global synaptic connectivity, highlighting the specificity and sensitivity of the system to detect microglial functional changes. This model captures genotype-dependent microglial reactivity (revealing phenotypes not fully captured by transcriptomic rofiling) and provides a physiologically relevant, tractable in vitro platform for dissecting microglial contributions to synaptic pathology in neurodegenerative disease.

## INTRODUCTION

Microglia are the resident immune cells of the central nervous system (CNS), contributing to homeostasis through synaptic pruning, phagocytosis of apoptotic cells, and immune surveillance [1–3]. Over the past century, our understanding of microglial biology has evolved from viewing them as passive bystanders to recognizing them as dynamic, multifunctional regulators of CNS homeostasis and pathology [4]. During brain development, microglia regulate synaptic connectivity by eliminating weak or redundant synapses via complement-dependent pruning [1,5]. In the adult brain, they continue to dynamically monitor and interact with synapses, modulating plasticity and responding to local activity [6]. Microglia share ontogeny and certain functional programs with other CNS-associated macrophages, such as perivascular and meningeal populations, which also contribute to neuroimmune surveillance and disease modulation [7]. In Alzheimer’s disease (AD), these physiological functions become dysregulated. Microglia become activated by amyloid-β (Aβ), shift toward a reactive phenotype, and can aberrantly engulf functional synapses, often through complement- or Fc receptor-mediated pathways, particularly involving the C1q–C3–CR3 (CD11b) axis [5,8,9]. These mechanisms are thought to contribute to early synaptic deficits and network dysfunction.

Genetic studies further support the central role of microglia in AD. Genome-wide association studies (GWAS) have identified multiple risk genes highly enriched in microglia, including TREM2, PLCG2, CD33, and ABI3, implicating immune pathways in disease susceptibility [10]. Despite this, in vitro models for studying microglial function often rely on neonatal, immortalized, or iPSC-derived cells that fail to replicate the transcriptomic and functional features of adult, disease-primed microglia [11,12]. Moreover, traditional co-culture systems rarely allow for compartmentalized manipulation or refined analysis of neuron–microglia interactions, particularly at the synaptic level, limiting mechanistic insight. Importantly, microglia derived from adult brain tissue retain transcriptional and epigenetic signatures that are often lost in neonatal or iPSC-derived models, underscoring their importance for disease-relevant functional studies [11].

To address this gap, we developed a compartmentalized microfluidic co-culture platform combining primary cortical neurons with adult microglia from WT and APP-transgenic mice. This system enables spatially controlled neuron–microglia interactions and quantitative assessment of microglial responses. As proof-of-concept, we applied lipopolysaccharide (LPS), a well-characterized proinflammatory stimulus, to elicit innate immune activation. LPS acts via TLR4/CD14 signaling, inducing cytokine release and morphological changes, pathways that overlap with neuroinflammatory mechanisms implicated in AD pathology [13,14]. This setup allowed us to compare morphological changes, cytokine release, and phagocytic behavior between genotypes, under controlled inflammatory conditions.

Transcriptomic profiling of isolated microglia was performed off-chip to identify genotype-specific inflammatory signatures, in line with recent single-cell studies describing a disease-associated microglia (DAM) phenotype, marked by upregulation of phagocytic and inflammatory genes in AD models [15].

Altogether, this approach allows for simultaneous evaluation of microglial morphology, cytokine release, synapse-specific phagocytosis, and transcriptional state in a controlled in vitro environment. We demonstrate that adult APP microglia exhibit heightened inflammatory and synapse-directed activity upon LPS stimulation, consistent with a disease-primed phenotype. This system offers a versatile and physiologically relevant in vitro platform to study genotype-specific microglial contributions to inflammation and synaptic dysfunction in neurodegeneration.

## MATERIALS AND METHODS

### 1. Animals and ethics

All experiments were done following the French ethical guidelines for studies on experimental animal project approval by the local ethics committee (CEEA75; APAFIS #32824-2021120518521661 v4). Mice were housed under SOPF conditions (22°C, 12 h light/dark cycle) with ad libitum access to food and water. Anesthesia was administered via intraperitoneal injection of ketamine (100 mg/kg) and xylazine (10 mg/kg), followed by transcardial perfusion with phosphate-buffered saline (PBS) containing 5 U/ml heparin to reduce blood contamination and improve microglial purity prior to tissue dissociation. APP-transgenic (hAPPJ20, PDGFAPPSw,Ind) [16] and T2A-tdTomato Hexb mice [12] mice were maintained on a C57BL/6J and C57BL/6N background respectively. To generate animals with endogenously labeled microglia, C57Bl/6N T2A-tdTomato Hexb mice were crossed with WT or APP animals. The resulting lines, referred to as To (WT background) and APPTo (APP background), carry endogenous, tdTomato-labeled microglia. All the experiments were done on F1 offspring 10-months-old mice.

### 2. Primary microglia isolation and culture

Adult microglia were isolated from the cortices of 10–11-month-old WT, APP, To and APPTo mice. Microglia isolation was performed as previously described [17,18], with minor modifications. Briefly, brains were dissected and cortices were enzymatically dissociated in Hibernate-A medium (Gibco) containing papain (20 U/mL), collagenase type IV (1 mg/mL), and DNase I (10 mg/mL; ca. 4000 Kunitz units/mL) at 37°C for 30 min with continuous gentle agitation. Every 10 min, tissue was gently triturated using a 1 mL pipette to promote dissociation. At the end of the incubation, the suspension was sequentially filtered through 100 µm, 70 µm, and 40 µm strainers to remove debris and undissociated tissue. Microglia were enriched by density gradient centrifugation [19] using a 16% OptiPrep solution (Sigma-Aldrich), centrifuged at 700 × *g* for 30 min at room temperature without brake, and collected from the interphase.

Cells were plated in six-well plates in plating medium (DMEM/F12 supplemented with 2% B27 (Miltenyi Biotec; with antioxidants), 1% GlutaMAX, 40 ng/mL FGF, and 2 μg/mL heparin) [20]. After 24 h, the medium was replaced with microglia maintenance medium (DMEM/F12, 10% fetal bovine serum [FBS], 1% GlutaMAX, 1% non-essential amino acids, 1% sodium pyruvate, 2 μg/mL HEPES, and 5 ng/mL macrophage colony-stimulating factor [M-CSF]). Cultures were maintained for 4–5 weeks with half-media changes twice per week to promote stabilization and expansion of viable microglia populations.

For each independent experiment, microglia were isolated from one WT and one APP adult mouse from the same litter whenever possible. Animals were generally matched for sex and age; however, due to constraints in colony availability, some experiments used non–sex-matched pairs. Cells were processed and cultured separately by genotype, and treatments (vehicle or LPS) were applied in parallel. Therefore, all independent experiments represent biological replicates from distinct animals, with multiple coverslips or devices per condition used as technical replicates.

### 3. Microfluidic devices

A tricompartmental neuron culture device was used [21]. Briefly, the device consists of a 450-μm-wide, 8.6-mm-long central chamber flanked by two 750-μm-wide, 4.5-mm-long side chambers. All three chambers are ca. 75 µm high. The left-side chamber (termed presynaptic) and the central chamber (termed synaptic) are interconnected via 4-µm-high, 450-µm-long parallel microchannels that narrow from an entry width of 10 μm to an exit width of 3 μm. The right-side chamber (termed postsynaptic) and the synaptic chamber are also interconnected via parallel microchannels with identical dimensions, except that they were 75 μm long.

Master patterns were fabricated at the IEMN (Lille, France) via two-step photolithography according to established procedures [22]. Ca. 4-mm-high polydimethysiloxane (PDMS) pads were replica molded. Access wells were punched at the termini of the central and side chambers using 3 mm and 4 mm biopsy punches (Harris Unicore), respectively. The devices were permanently bonded to 24 mm × 50 mm glass coverslips (Menzel) via O_2_ plasma (Diener, Ebhausen, Germany). Devices were sterilized under UV light (Light Progress, Anghiari, Italy) for 30 min, then coated with poly-D-lysine (PDL; 0.1 mg/mL in PBS) overnight at 4°C. After rinsing with PBS, a second coating with laminin (20 μg/mL in PBS) was applied overnight at 4°C. Before cell seeding, devices were acclimated at 37°C for 30 min.

### 4. Primary neuronal culture

Primary cortical neurons were prepared from embryonic day 14–15 (E14–E15) wild-type C57BL/6N mouse embryos. Pregnant females were euthanized by CO_2_ inhalation followed by cervical dislocation, and embryos were rapidly collected in cold dissection buffer (HBSS supplemented with 10 mM HEPES and 1% Penicillin/Streptomycin). Cortices were isolated under a stereomicroscope, and meninges were carefully removed. Tissue was digested for 10 min at 37°C in 0.25% trypsin-EDTA (Gibco) and DNase I (100 μg/mL), followed by gentle mechanical dissociation. Cells were counted and plated into the coated devices (50,000 neurons in each pre- and postsynaptic chamber) in Neurobasal Plus medium supplemented with 2% B27 Plus and 1% GlutaMAX. Neurons were maintained in culture for 14-16 days, with half-media changes performed every 3–4 days. A lentiviral construct encoding a PSD95-targeting EGFP probe under the human Synapsin I promoter (VectorBuilder, VB220701-1052zbm) was used to visualize synaptic material. This construct is based on a recombinant intrabody strategy targeting endogenous PSD95 [23] and was adapted for neuron-specific expression. Cortical neurons were transduced at day in vitro 7 (DIV7) and subsequently co-cultured with microglia at DIV9 (Figure 1).

**Figure 1.**
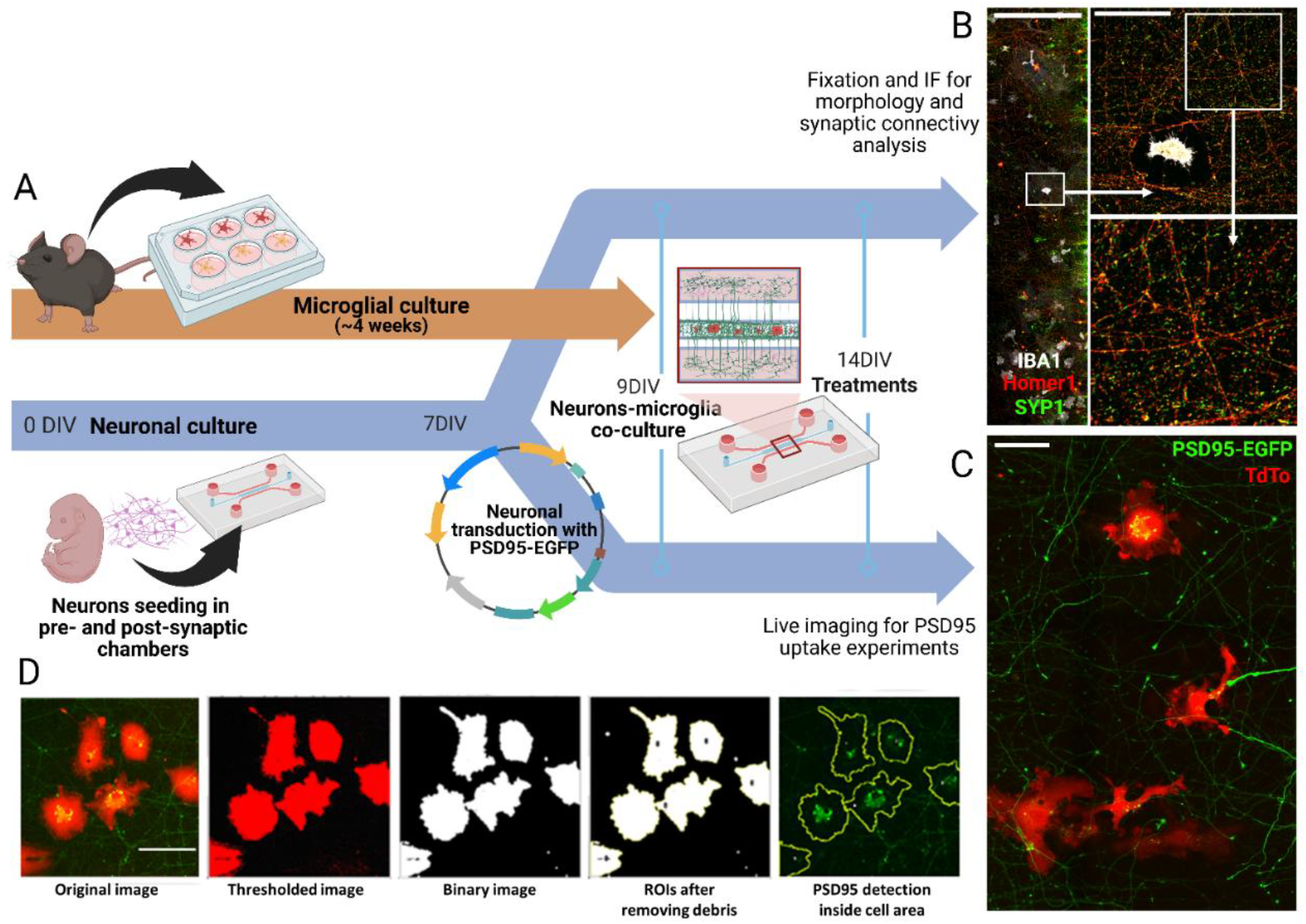
Workflow of the microfluidic co-culture system for microglial-synaptic interaction analysis. **(A)** Schematic timeline of the co-culture model. Adult microglia were isolated and expanded in culture for ca. 4 weeks. In parallel, primary cortical neurons from E14–E15 embryos were cultured in microfluidic devices, seeded into pre- and postsynaptic compartments at DIV0. At DIV7, neurons destined for live imaging experiments were transduced with a lentiviral construct encoding a PSD95-targeting EGFP probe. At DIV9, adult microglia were added to the synaptic chamber (ca. 100 cells/device), initiating the co-culture phase. Scale bars = 500 µm (left) and 50 µm (right). **(B)** Immunocytochemistry and morphological/synaptic analysis: Devices were fixed at DIV14 and immunostained for IBA1 (microglial marker), Synaptophysin (SYP; presynaptic marker), and Homer1 (postsynaptic marker) to assess microglial morphology and synaptic connectivity. Representative images show IBA1^+^ microglia in proximity to mature SYP–Homer1 pairs in the synaptic chamber. Scale bar = 50 µm. **(C)** Live imaging: At DIV14–15, devices were imaged live to quantify synaptic material internalization. Representative image shows tdTomato-expressing microglia interacting with PSD95-EGFP neuronal processes. Scale bar = 50 µm. **(D)** Quantitative analysis of PSD95 uptake: IBA1 and tdTomato signals were segmented to generate binary masks and regions of interest (ROIs), used to assess morphology and quantify internalized PSD95-EGFP signal per cell. Image processing included thresholding and morphological operations. Created in BioRender.com.

### 5. Microglia/neurons co-culture

At DIV9, microglia previously maintained in 6-well plates were harvested and seeded into microfluidic co-culture devices containing primary cortical neurons, as described in Figure 1. Microglia detachment was performed in two sequential steps: first, cells were incubated on a rocker with 50 U/mL dispase (5 min, 37°C, 180 rpm), followed by 0.05% trypsin-EDTA (5 min, 37°C, 180 rpm). Detached cells were centrifuged at 300 × *g* for 5 min, resuspended in co-culture medium (Neurobasal supplemented with 2% B27, 5% FBS, and 0.25% GlutaMAX), and counted.

Microglia were maintained for 3–4 weeks in 6-well plates, typically using two wells per genotype. At the end of this period, total yields ranged from ca. 100,000 to 600,000 cells per genotype, with APP lines generally yielding higher cell numbers than WT, indicating a slightly faster proliferation under our culture conditions. For microglia seeding, neuronal medium was removed from all chambers of the device. This step was required to transition to co-culture medium and also helped minimize fluid pressure that could displace microglial cells during seeding. Approximately 100 microglial cells were added to the synaptic chamber. After allowing the cells to settle, the entire device was gently filled with fresh co-culture medium to support both cell types and was maintained in a humidified incubator (37°C, 5% CO_2_). Neurons continued to mature within the devices alongside microglia until DIV14, when functional assays were performed.

### 6. Experimental treatments

At DIV14, co-cultures were subjected to inflammatory stimulation and complement receptor blockade. To ensure an even distribution of the treatment and preserve the conditioned microenvironment, medium from the synaptic chamber was first collected and mixed with the appropriate reagent (e.g., LPS or anti-CD11b antibody) in a microcentrifuge tube. The treatment mixture was then gently reintroduced into the synaptic chamber. LPS (100 ng/mL; Sigma-Aldrich) was applied for 6 h, followed by a complete medium replacement and a 16–18 h recovery period in co-culture medium. To block CD11b, the anti-CD11b monoclonal antibody (clone M1/70.15; 10 μg/mL; Bio-Rad, MCA74EL) was added 24 h prior to LPS treatment and maintained throughout the stimulation and recovery phases. Imaging was performed at baseline, immediately after treatment, and after the recovery period, when applicable.

General phagocytic activity of primary microglia from adult mice was assessed using pHrodo™ Green Zymosan particles (Thermo Fisher Scientific, P35365). Briefly, at DIV14, co-cultures were treated with vehicle or LPS, with or without pre-treatment with anti-CD11b monoclonal antibody as described above. 3 h after the start of LPS treatment, pHrodo™ Zymosan particles (prepared according to the supplier’s protocol) were added to the synaptic chamber at the recommended working concentration. Cells were maintained at 37°C and 5% CO_2_ until imaging.

### 7. Immunocytochemistry

At the end of the experimental protocol, i.e., 16–18 h after treatment, media was removed from all chambers and cultures were fixed with 4% paraformaldehyde (PFA) in PBS for 15 min at room temperature. After fixation, cells were rinsed three times with PBS, permeabilized with 0.3% Triton X-100 in PBS for 10 min, and blocked overnight at 4°C with blocking buffer containing 1% bovine serum albumin (BSA), 0.1% Tween-20, and 5% donkey serum in PBS. Primary antibodies were applied overnight at 4°C in the same buffer. The following primary antibodies were used: rabbit anti-Iba1 (1:1000, FUJIFILM Wako), mouse anti-Tau (1:400, Synaptic Systems), goat anti-Homer1 (1:250, Synaptic Systems), and chicken anti-synaptophysin (1:500, Synaptic Systems). After washing, secondary antibodies were applied for 1 h at room temperature. The following secondary antibodies were used: donkey anti-rabbit Alexa Fluor 647 (1:500), donkey anti-mouse DyLight 405 (1:500), donkey anti-goat Alexa Fluor 555 (1:500), and donkey anti-chicken Alexa Fluor 488 (1:500). Samples were mounted using 90% glycerol in PBS and stored at 4°C until imaging (Figure 1B).

### 8. Image acquisition and analysis

#### A. Cell morphology and PSD95 phagocytosis

Immunostained microfluidic devices were imaged using a Spinning Disk X-Light V3 microscope (Nikon) equipped with a 40× objective. A *z*-step of 0.3 μm was used to obtain stacks. Mosaic images covering the entire synaptic chamber were acquired for each device. Image analysis was performed using custom ImageJ (NIH; RRID:SCR_003070) macros developed for microglial morphometry and synaptic uptake quantification.

For morphometric analysis, microglial cells were segmented based on Iba1 fluorescence. Background noise was reduced using the despeckle tool, and binary masks were generated by automated thresholding followed by binary morphological operations (fill holes, dilate, erode) to refine cell contours (see Figure 1D). Cells at image borders were excluded. A minimum area threshold of 562 μm^2^ was applied to exclude debris, corresponding to the first quartile of microglial area dataset for the WT+Vehicle condition, obtained across four independent experiments.

To quantify PSD95 uptake by microglia, corrected GFP signal intensity per cell was computed as (Mean Intensity – Mean Background) × Area (Figure 1C). Microglial masks generated from tdTomato fluorescence (as described above) were used to define the regions of interest (ROIs) for signal extraction. Background values were calculated per coverslip using non-cell regions. For each condition, the median corrected intensity across all segmented microglial cells from a single coverslip was used as the statistical unit. Each coverslip was treated as an independent biological replicate, and group-level comparisons were performed based on the mean ± SEM of these per-coverslip medians. A total of 400–650 cells per condition were analyzed, derived from 8–9 coverslips across 5 independent experiments.

Morphology data violated the assumption of homogeneity of variances (Brown–Forsythe test); group comparisons were thus performed using the Kruskal–Wallis test with Dunn’s post hoc correction. In contrast, PSD95 phagocytosis data met the assumptions for parametric testing and were analyzed using one-way ANOVA followed by Tukey’s multiple comparisons test.

#### B. Synaptic connectivity

For quantification of synaptic connectivity, high-resolution z-stacks of the synaptic chamber were acquired using a 60× oil-immersion objective (NA 1.4) with a *z*-step of 0.2 μm, on a spinning disk confocal system (X-Light V3, CrestOptics). Image stacks were deconvolved using Huygens Professional software (Scientific Volume Imaging; RRID:SCR_014237), and 3D puncta of Synaptophysin (SYP) and Homer1 were reconstructed in Imaris (Bitplane, Zürich; RRID:SCR_007370). Synaptic connectivity was calculated as previously described [21] by identifying apposed SYP– Homer1 pairs using a custom, MATLAB-based spatial colocalization algorithm. This method relies on the spatial proximity of pre- and postsynaptic puncta as a proxy for functional synapses, enabling the detection of net changes in mature synapse number.

#### C. pHrodo™ Zymosan uptake

Live imaging was performed 2 h after particle addition. Internalized pHrodo™ Zymosan signal was quantified per tdTomato^+^ microglial cell using the same segmentation and background subtraction pipeline described above. For each condition, the median corrected intensity across all segmented cells on a coverslip was used as the statistical unit.

### 9. IL-1β quantification

IL-1β secretion was measured in primary adult microglia cultured on 13 mm glass coverslips. Microglia were seeded following the same protocol as for microfluidic co-cultures, at a density of 20,000 cells per coverslip. After 4–5 weeks in culture, cells were treated with 100 ng/mL LPS for 6 h in complete microglia medium. Media was then replaced with fresh medium, and cells were allowed to recover for 16 h. At the end of the recovery period, 500 μL of medium was collected from each well and concentrated using 5 kDa centrifugal filters (Vivaspin 500, Sartorius) at 10,000 × *g* for 30 min. IL-1β levels were quantified using the AlphaLISA immunoassay (PerkinElmer, AL503C/F) according to the manufacturer’s instructions. Data are expressed as Alpha counts.

Eight samples per condition were analyzed, each derived from an independent experiment (i.e., different animals). Only samples showing at least a two-fold increase compared to their vehicle controls and no outlier behavior (based on the 1.5× interquartile range method) were included. All values were normalized to the mean of the WT+vehicle group. Statistical analysis was performed using one-way ANOVA followed by Tukey’s post hoc test. Data met assumptions for parametric testing, including normality and homogeneity of variance.

### 10. Transcriptomic analysis

Adult microglia were cultured for four weeks as described previously, then detached and seeded onto 24-well plates containing 13 mm glass coverslips (25,000 cells per well) to replicate the substrate properties of microfluidic devices. Four days later, cells were treated with either vehicle or LPS (100 ng/mL) for 6 h. Media was then replaced with fresh medium, and cells were allowed to recover for 16 h before RNA extraction. Total RNA was isolated using the RNeasy Plus Mini Kit (Qiagen, #74136) according to the manufacturer’s protocol. Only samples with RNA integrity number (RIN) > 8.5 were considered for sequencing. A total of 19 samples were processed: WT+Vehicle (n = 4), WT+LPS (n = 5), APP+Vehicle (n = 5), and APP+LPS (n = 5).

Libraries were prepared using the QuantSeq 3′ mRNA-Seq Library Prep Kit FWD (Lexogen), which enriches for 3′ ends of polyadenylated transcripts. External RNA Controls Consortium (ERCC) spike-in mixes were included to assess detection sensitivity. Unique Molecular Identifiers (UMIs) were attached prior to PCR amplification to reduce biases and improve quantification accuracy. Sequencing was performed on an Illumina NovaSeq 6000 platform with single-end 100 bp reads.

Raw reads were quality-checked using FastQC [24] and trimmed with Trimmomatic [25] to remove adapters and low-quality bases. Pseudo-alignment was performed using Kallisto [26] against the Ensembl mouse transcriptome [27]. Transcript abundance estimates were imported using the tximport package in R [28], and differential expression analysis was conducted with DESeq2 [29]. Genes with total counts < 100 were excluded prior to normalization. Differentially expressed genes (DEGs) were defined as those with |log_2_ fold change| > 1 and a false discovery rate (FDR) < 0.05.

### 11. Statistical analysis

Statistical analyses were performed using GraphPad Prism 10. Normality and variance homogeneity were assessed using Shapiro–Wilk and Brown–Forsythe tests, respectively. Data were analyzed using one-way ANOVA or the Kruskal–Wallis test, as appropriate. Outliers were excluded using the 1.5× IQR method applied consistently across datasets. Results are presented as mean ± SEM unless otherwise indicated, and significance level was defined as *p* < 0.05.

## RESULTS

### APP microglia exhibit a primed inflammatory state and respond robustly to LPS

To validate that our co-culture system preserves microglial immunocompetence, we quantified IL-1β levels in culture supernatants 16 h after LPS treatment using AlphaLISA. LPS stimulation induced a significant increase in IL-1β secretion in both WT and APP microglia (ANOVA p < 0.0001), with fold changes of 2.2 in WT (p < 0.0001) and 2.9 in APP (p < 0.0001). Moreover, APP microglia released significantly more IL-1β than WT following LPS (FC = 1.3, p = 0.0102), indicating a hyperresponsive inflammatory phenotype (Figure 2A). Baseline IL-1β levels were comparable between genotypes under vehicle conditions. These findings support the presence of a primed proinflammatory state in APP microglia, consistent with the concept of microglial sensitization in neurodegeneration, where phagocytosis-driven secretomes can actively modulate neurogenesis in vivo [30].

**Figure 2.**
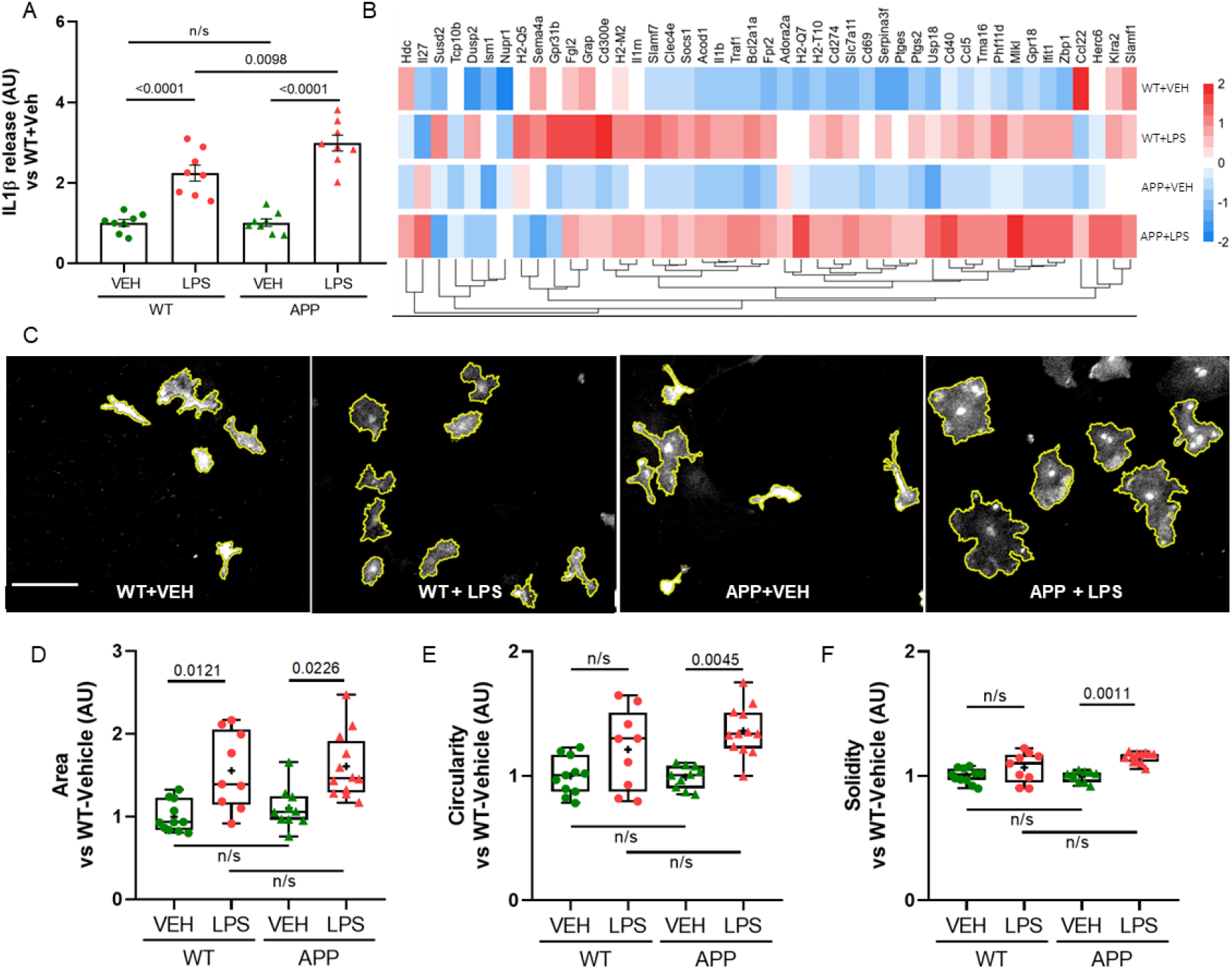
Validation of the microfluidic model and differential microglial activation. **(A)** IL-1β levels were measured in culture media 16 h after LPS (100 ng/mL) or vehicle treatment using AlphaLISA. LPS induced a robust increase in IL-1β secretion in both WT and APP microglia, with APP showing significantly higher levels than WT after stimulation. Data represent mean ± SEM; *n* = 8 per group (each from separate WT and APP mice). One-way ANOVA with Tukey’s post hoc test. **(B)** Heatmap showing z-score normalized expression of selected inflammation-related genes from bulk RNA-seq, including cytokines (*Il1b, Tnf, Ccl3*), interferon-responsive genes (*Usp18, Ifit1, Herc6, Zbp1*), prostaglandin synthesis (*Ptgs2*), and MHC-related genes (*H2-Q7, H2-M2, H2-T10*). LPS broadly upregulated these transcripts, with stronger induction in APP microglia, indicating heightened inflammatory responsiveness. **(C)** Representative images of IBA1^+^ WT and APP microglia after 6 h treatment with vehicle or LPS. Yellow outlines indicate segmented cell bodies used for morphometric analysis. Scale bar = 100 µm. **(D–F)** Quantification of shape descriptors in WT and APP microglia, normalized to WT+Vehicle: area (**D**); circularity (**E**), and solidity (**F**). LPS significantly increased area in both genotypes and increased circularity and solidity in APP microglia only, indicating a shift toward a reactive, less ramified morphology. Each data point represents the median value across all segmented microglia from a single device. Group comparisons are based on the mean ± SEM of these per-device medians. n = 9–12 devices from ≥4 independent experiments (each from separate WT and APP mice). Statistical test: Kruskal–Wallis with Dunn’s post hoc correction; adjusted p-values shown.

To further characterize this inflammatory response, we performed bulk RNA-seq on microglia exposed to LPS or vehicle. As a quality control step, we verified the expression of canonical microglial markers in the raw RNA-seq data. Canonical microglial markers were consistently expressed across all samples, including homeostatic genes such as *Cx3cr1, Tmem119*, and *P2ry12*, as well as the pan-microglial identity gene *Hexb*, which showed particularly high abundance (>10,000 normalized counts). Additional microglial signature genes, including *Fcrls, Sall1, Siglech*, and *Gpr34*, were also detected, supporting the purity of the microglial cultures and confirming that the transcriptomic profiles derived from genuine microglial populations. Genes associated with cytokine signaling (*Il1b, Tnf, Ccl3*), interferon response (*Ifit1, Zbp1, Usp18*), prostaglandin synthesis (*Ptgs2*), and antigen presentation (*H2-Q7, Cd74*) were broadly upregulated upon LPS exposure in both genotypes, with stronger induction in APP microglia (Figure 2B). Although only a subset of genes reached statistical significance, the global patterns were consistent with a primed immune phenotype.

### LPS induces morphological activation in adult microglia

Adult microglia from WT and APP transgenic mice exhibited robust morphological responses to LPS stimulation. Both WT and APP microglia showed a significant increase in cell area after LPS treatment compared to vehicle controls (Kruskal–Wallis test; p = 0.0002; WT+LPS vs. WT adj. p = 0.0121; APP+LPS vs. APP adj. p = 0.0226). APP microglia also displayed increased circularity (p = 0.0045) and solidity (p = 0.0011), indicative of a rounder, less ramified morphology, consistent with activation. These findings suggest that adult APP microglia undergo a morphological shift in response to LPS, consistent with a reactive phenotype. However, morphology alone may not fully capture functionally distinct activation states.

### APP microglia exhibit increased PSD95 phagocytosis after LPS stimulation without altering global synaptic connectivity

To assess microglial interaction with synapses, neurons were transduced at DIV7 with a lentivirus encoding a GFP-tagged nanobody that binds endogenous PSD95. Microglia were seeded into the synaptic chamber 48 h after neuronal transduction. Live imaging was performed at baseline and again at 16 h after treatment with LPS (100 ng/mL) or vehicle (Figure 3A). Internalized PSD95 signal was quantified per cell and aggregated at the coverslip level. Under basal conditions, APP microglia showed an average ∼40% higher PSD95 intensity compared to WT, although this difference was not statistically significant, it could indicate a tendency toward greater constitutive uptake of synaptic material. Following LPS stimulation, APP microglia showed a significant increase in PSD95 internalization (p = 0.0092), while WT microglia displayed a mild, non-significant increase (Figure 3B). The LPS-induced response was significantly greater in APP compared to WT cells (p = 0.0016), indicating a heightened synapse-engulfing capacity in the disease model.

**Figure 3.**
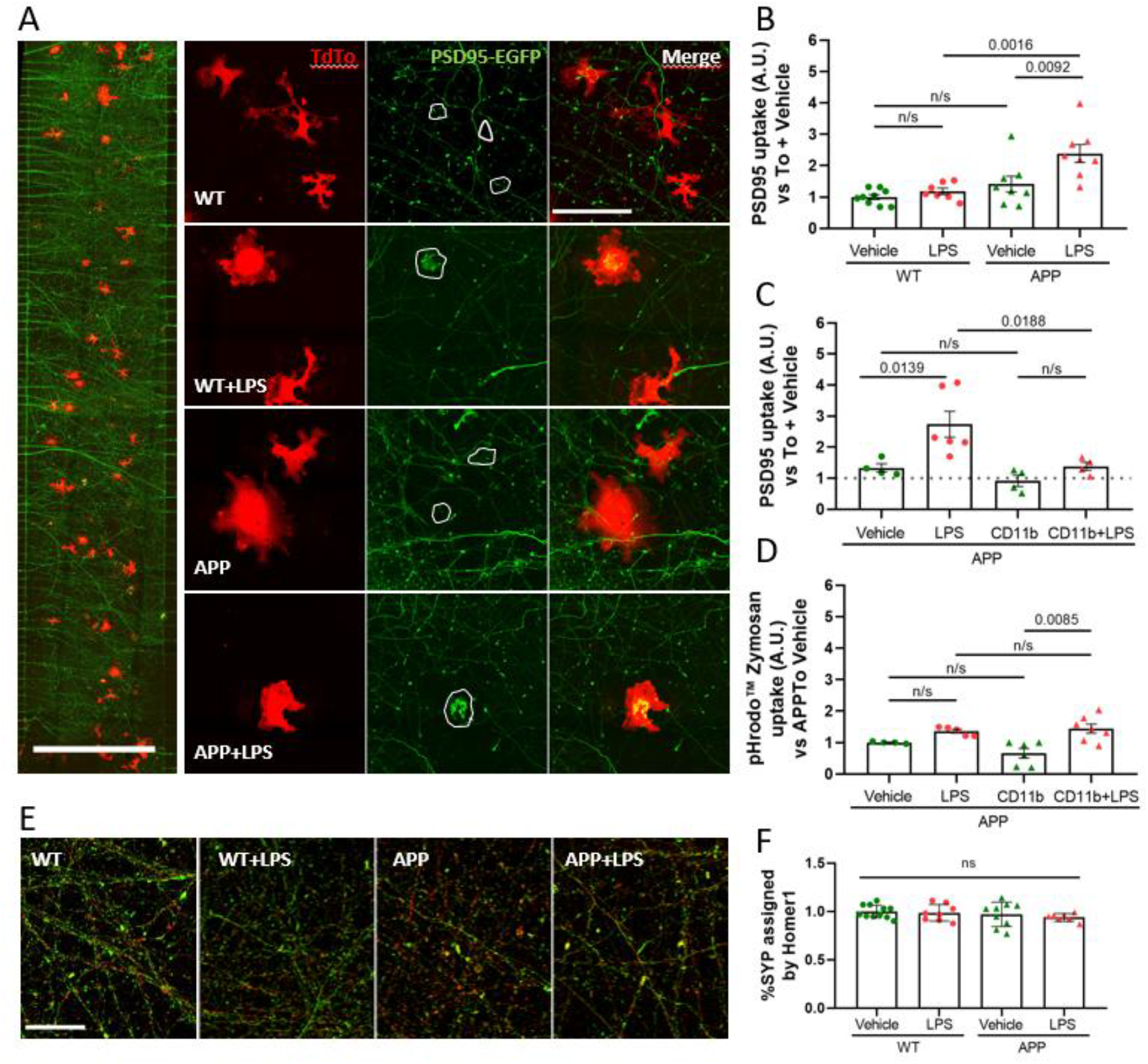
Selective enhancement of synaptic phagocytosis in APP microglia. **(A)** Representative mosaic image from the live PSD95 uptake assay (left). Images show WT and APP microglia under vehicle or LPS (100 ng/mL). TdTomato^+^ microglia (red) interact with PSD95-EGFP^+^ neuronal structures (green). White outlines indicate microglial ROIs used for quantification. **(B)** Quantification of PSD95 internalization per microglial cell. LPS selectively enhanced PSD95 uptake in APP microglia, suggesting increased synaptic targeting. One-way ANOVA with Tukey’s post hoc test. **(C)** CD11b blockade reduced PSD95 internalization in LPS-treated APP microglia, supporting a partial dependence on CD11b signaling. **(D)** Uptake of pHrodo™ Green Zymosan, a non-specific substrate, remained unchanged, indicating that the phagocytic increase was specific to synaptic material. **(E)** Representative immunostaining of synaptic compartments shows Synaptophysin (SYP, green) and Homer1 (red) puncta. **(F)** Quantification of synaptic connectivity, defined as the proportion of SYP puncta assigned by Homer1. No differences were observed between conditions, suggesting that LPS did not disrupt overall synaptic architecture. To avoid local effects of microglial contact, analysis was restricted to regions ≥75 μm from the nearest microglial soma. Data in (B–D, F) represent mean ± SEM, normalized to vehicle of each genotype; *n* = 4–6 coverslips per condition from ≥3 independent experiments. Scale bars: (A) 500 µm (left), 100 µm (right); (E) 20 µm.

To evaluate synaptic integrity under inflammatory challenge, we quantified synaptic connectivity based on apposed SYP– Homer1 puncta within the synaptic chamber as described in Methods. To avoid confounding effects of direct microglia–synapse contact, synaptic connectivity was quantified in areas located at least 75 μm away from the nearest microglial cell. This spatial buffer was applied to exclude regions potentially affected by local engulfment or remodeling activity. Despite robust microglial activation and increased PSD95 uptake, LPS treatment did not significantly alter synaptic connectivity in either genotype (Figure 3E-F). These results suggest that, over short timescales, microglial secretory activity alone is not sufficient to cause a measurable synaptic loss, highlighting the importance of direct contact or prolonged interaction for structural synaptic changes.

### Increase in synaptic phagocytosis in LPS-stimulated APP microglia depends on the complement receptor CD11b

To explore potential mechanisms of synaptic phagocytosis in APP microglia, we assessed the role of complement receptor CD11b. Blocking CD11b with a specific antibody (clone M1/70.15; BioRad) did not affect PSD95 internalization in non-stimulated APP microglia but reduced the LPS-induced PSD95 uptake to control levels (Figure 3C), confirming the involvement of the C1q/C3–CR3 pathway in synaptic phagocytosis [5]. To assess bulk, non-specific phagocytic activity, we used pHrodo™ Zymosan, a pH-sensitive fluorescently labeled yeast particle commonly employed to monitor general phagocytosis in microglia. In contrast to the PSD95 results, LPS stimulation did not significantly increase pHrodo™ Zymosan uptake compared to vehicle, suggesting that APP microglia do not exhibit a strong enhancement of bulk phagocytic capacity under inflammatory conditions. However, CD11b blockade alone reduced pHrodo™ Zymosan uptake, and this reduction was partially reversed by LPS co-treatment (p = 0.0085; Figure 3D). Notably, LPS also resulted in significantly higher uptake compared to CD11b blockade alone (p = 0.0180), although levels remained comparable to baseline. These findings suggest that while LPS can engage CD11b-independent pathways to support bulk phagocytosis, this response is limited in magnitude. In contrast, synaptic phagocytosis showed a robust, CD11b-dependent enhancement under the same conditions. This highlights a mechanistic divergence between synaptic and non-specific engulfment and supports the notion that synapse-specific phagocytosis is more selectively regulated and amplified in APP microglia during inflammatory challenge.

## DISCUSSION

Understanding how genetic risk for AD shapes microglial behavior remains a critical challenge. In this study, we present a microfluidic co-culture system that enables adult microglia from WT and APP-transgenic mice interact with synapses of primary cortical neurons. Using this platform, we demonstrate that APP microglia display exaggerated morphological changes, elevated IL-1β release, and increased uptake of synaptic markers in response to LPS stimulation. These findings support the concept of a primed or hyperresponsive state in disease-associated microglia. Importantly, the microfluidic architecture not only allows for controlled co-culture of adult microglia and neurons but also permits spatial segregation of neuronal somata from the synaptic network. This configuration enables targeted stimulation of synapses, localized microglia–synapse interactions and refined morphometric analyses. Additionally, the system requires low cell numbers and reagent volumes, making it suitable for scarce or precious samples and scalable for pharmacological testing.

### Genotype-specific microglial activation and inflammatory priming

IL-1β is a key proinflammatory cytokine involved in microglial neurotoxicity and synaptic dysfunction [13,31–33]. In our model, APP microglia secreted significantly more IL-1β than WT microglia following LPS stimulation, despite comparable baseline cytokine levels. These results are consistent with the concept of microglial “priming” (a state characterized by exaggerated responses to secondary insults) even in the absence of overt activation at rest. This phenomenon has been well-documented in aging models and neurodegenerative conditions [34] and is thought to underlie chronic innate immune activation observed in Alzheimer’s disease and related disorders [35]. More recent studies have shown that primed microglia exhibit transcriptionally distinct and epigenetically biased profiles favoring proinflammatory responses [15,36,37]. Importantly, pathways linked to TREM2 and APOE have been implicated in modulating this sensitized state, thereby connecting AD genetic risk to immune reactivity [15,38]. These findings support our interpretation that the APP genotype confers a primed state at baseline, which becomes apparent under exogenous stimulation. Such genotype-driven differences in microglial responsiveness align with recent comprehensive reviews highlighting the diversity of microglial phenotypes, their developmental origins, and their context-dependent roles in neurodegenerative disease [4,7]. While our model does not capture long-term environmental influences, it reflects genotype-intrinsic features of immune programming.

Microglial morphological activation, characterized by increased cell area and loss of ramification, is a hallmark of inflammatory response [39–41]. In our system, both WT and APP microglia underwent significant morphological changes after LPS treatment. However, APP cells exhibited a more pronounced shift in circularity and solidity, consistent with a transition to an amoeboid phenotype. These findings suggest that APP microglia are more sensitive to proinflammatory stimuli. This observation aligns with previous in vivo studies reporting heightened microglial reactivity in APP mice [42,43], and demonstrates that our in vitro platform can capture genotype-specific immune responses using adult microglia.

### Synapse-specific phagocytosis and functional specificity

Whether inflammatory stimuli globally enhance microglial phagocytosis or selectively increase uptake of synaptic substrates remains an open question in the field. Our experimental design, which includes parallel assessment of synapse-targeted and non-specific engulfment, enables direct comparison of substrate specificity under controlled conditions.

To assess synapse-specific phagocytosis, we used a PSD95-GFP nanobody system and observed that APP microglia internalized significantly more PSD95 after LPS stimulation compared to WT. While both genotypes secreted IL-1β upon LPS exposure, only APP microglia showed robust synaptic uptake. This divergence suggests that, although classical inflammatory pathways, such as TLR4-driven cytokine release, are broadly inducible, synaptic phagocytosis operates under a more selective, threshold-dependent mechanism.

A trend toward elevated PSD95 uptake in APP microglia under basal conditions could indicate a chronically active or dysregulated surveillance state, in line with reports of early synaptic loss mediated by microglia in AD models [44,45]. Notably, blocking CD11b effectively abolished the LPS-induced increase in PSD95 uptake by APP microglia (Figure 5C), confirming the role of complement signaling in synapse-specific phagocytosis. However, this blockade had no effect under basal conditions, implying that additional, non-complement pathways may mediate constitutive synaptic uptake.

Despite robust microglial activation and increased PSD95 uptake, we did not detect significant changes in the global synaptic connectivity in this paradigm. This may reflect the short duration of exposure or the resilience of mature neuronal networks in vitro. Importantly, synaptic connectivity was quantified at a distance from microglial surfaces to avoid regions with obvious local disruptions or neuronal network loss. As such, this approach captures the broader, secreted effects of microglia on the synaptic network rather than direct cell-contact–mediated pruning. It remains possible that focal synaptic elimination occurred in close proximity to microglial cells, but its effect was diluted in global connectivity analyses. Nevertheless, the ability to quantify synaptic integrity via high-resolution imaging remains a key advantage of this system.

In contrast to synaptic uptake, exposure to pHrodo™ Zymosan, a generic phagocytic substrate, did not reveal significant differences across genotypes or treatments. Neither LPS nor CD11b antibody significantly altered pHrodo™ Zymosan uptake, suggesting that general phagocytosis is less sensitive to genotype or inflammatory context. These findings reinforce that synaptic engulfment is a distinct, regulated process requiring specific cellular cues, such as neuronal distress or opsonization. Our comparison between PSD95 and pHrodo™ Zymosan uptake highlights the specificity of microglial responses and the limitations of general phagocytic assays in modeling disease-relevant activity. This interpretation is supported by findings showing that microglia respond distinctly depending on the phagocytic substrate they interact with, and that their secretory phenotype is shaped by the nature of the engulfed material, highlighting the importance of using target-specific assays to reveal functional heterogeneity[30]. Thus, our system is particularly well-suited to dissect microglial responses to synaptic pathology, rather than their general phagocytic activation.

### Advantages and validation of the microfluidic co-culture model

This study demonstrates the utility of a compartmentalized microfluidic system for dissecting microglial functions in a controlled, neuron-rich environment. By physically separating neuronal somata from their neurites and synapses, the model allows one to study microglia–synapse interactions independently of neuronal somata and to use high-resolution imaging of synaptic structures. This spatial organization not only enhances biological relevance but also supports more refined imaging analyses than traditional well-based co-cultures. Furthermore, this model requires low cell and reagent volumes, making it well-suited for experiments using scarce or age-restricted material such as adult microglia, and overcoming the low-yield limitations reported for conventional adult microglial cultures [18,44]. Its modularity and compatibility with live imaging, genetic labeling, and localized treatments (e.g., LPS, Aβ) enable flexible experimental designs tailored to address diverse mechanistic questions. However, some limitations remain. While robust for functional readouts such as phagocytosis and morphological analysis, the small number of microglia seeded per device currently precludes direct transcriptomic analysis from within the chip. Instead, gene expression profiling must be performed on parallel bulk cultures under matched conditions, potentially limiting direct comparability.

While transcriptomic profiling was not the main focus of this study, we performed bulk RNA-seq as an exploratory tool to characterize the inflammatory response of adult microglia in culture. As detailed in the supplementary data, canonical microglial markers were well preserved, and several inflammation-related genes (such as *Il1b, Tnf, Ccl3*, and *Dhx33*) were differentially expressed following LPS stimulation. Although most of the observed effects were moderate in magnitude, they aligned well with the functional profile of APP microglia. Notably, prolonged in vitro maintenance prior to co-culture (21–28 days) has been shown to alter the homeostatic transcriptomic identity of ex vivo microglia [11,45] potentially constraining the dynamic range of transcriptional responses.

Nevertheless, key functional properties (such as cytokine release and substrate-specific phagocytosis) can be modulated or re-engaged upon stimulation, even in long-term cultures [11,45]. This may explain why APP microglia in our system exhibited a robust and selective phagocytic response despite partial loss of homeostatic gene signatures. In this context, the transcriptomic data support the overall interpretation of an exaggerated inflammatory phenotype in APP microglia and are consistent with our functional findings.

### Limitations and future applications of the in vitro model

While the aim of this study was to validate a functional in vitro model to assess microglial genotype-specific responses, several limitations should be acknowledged. The use of a single proinflammatory stimulus (LPS) provided a well-controlled, reproducible context to reveal differences between WT and APP microglia; however, LPS does not fully mimic endogenous stimuli relevant to neurodegeneration. Oligomeric Aβ species, for example, engage distinct receptor pathways (including CD36, CD14, integrins, and multiple TLRs) and elicit sustained, context-dependent responses related to synaptic dysfunction and neurotoxicity [46,47]. Unlike the acute and robust activation induced by LPS, Aβ species often elicit more gradual and sustained responses, modulating microglial phagocytosis in an aggregation-dependent manner and inducing transcriptional programs associated with DAM phenotypes [46]. Early foundational studies by Barres and colleagues demonstrated that glial cells, including microglia, actively regulate synaptic connectivity via complement-dependent pruning during development, establishing a paradigm for immune-mediated synapse refinement [5].

Furthermore, microglia are not static surveying cells: Tremblay and co-workers showed that microglial processes dynamically contact and monitor synaptic elements in the healthy brain, modulating with experience even in the absence of pathology [6]. This dynamic interaction highlights the utility of our live-cell platform to explore subtler, experience-dependent microglial behaviors.

Despite these differences, LPS was intentionally selected as a proof-of-concept trigger due to its robust and rapid effects. Importantly, our microfluidic co-culture system remains fully compatible with localized delivery of Aβ, tau, or other CNS-derived signals into the synaptic compartment, enabling precise interrogation of microglial responses to physiologically more relevant cues. This platform therefore provides a valuable tool to detect genotype-specific reactivity and functional differences in synapse-targeted phagocytosis. Previous studies have shown that microglial responses to Aβ differ from those to LPS in terms of cytokine secretion and synaptic engagement [48], and future work applying this platform to pathological ligands may uncover distinct or complementary immune phenotypes.

From a design standpoint, each independent experiment in our study used microglia isolated from distinct adult WT and APP mice. Animals were generally matched for age and sex, although sex-matching was not possible in all cases due to colony constraints. Reported n-values thus represent biological replicates, with multiple coverslips or devices analyzed per condition as technical replicates. Expanding the number of biological replicates and explicitly incorporating sex as a variable in future experiments would enhance the statistical resolution and translational relevance of this model.

Adult microglia were maintained in vitro for 21–28 days prior to co-culture, a step necessary for stabilization and reproducibility. However, long-term culture has been shown to alter the homeostatic transcriptional identity of microglia [45], which may constrain basal gene expression or responsiveness in some assays.

Prior studies have used ex vivo preparations with adult microglia, such as acute spinal cord slices for injury responses [49], or in vitro cultures of adult microglia on standard plates to assess phagocytosis and cytokine release [18,50,51]. Others have implemented microfluidic systems with microglial cell lines to study calcium dynamics [52]. To our knowledge, no existing approach combines *in vitro*–cultured adult brain microglia with spatially compartmentalized microfluidics, enabling simultaneous synapse-specific phagocytosis assays, high-content morphological analysis, and matched transcriptomic profiling. Moreover, this platform supports the functional interrogation of genotype-specific phenotypes in disease-relevant microglial populations. Taken together, it bridges a critical methodological gap between in vivo complexity and in vitro precision.

## CONCLUSION

This study introduces and functionally validates a novel microfluidic co-culture system combining adult microglia from APP-transgenic mice with mature cortical neurons harvested from embryonic mice. Unlike traditional neonatal or immortalized models, this platform preserves adult microglial features and enables spatially resolved neuron–microglia interactions. Upon LPS stimulation, APP microglia exhibited heightened inflammatory and synaptic responses, including increased IL-1β release, morphological activation, and enhanced phagocytosis of endogenous PSD95 with no alteration of general phagocytic behavior. These findings reveal genotype-dependent reactivity that is not fully captured by transcriptomic profiling. Notably, this platform also allows for assessment of global synaptic connectivity, although no differences were observed under the current experimental conditions.

Rather than dissecting specific pruning pathways, our aim was to establish a robust and tractable system bridging in vivo relevance and in vitro control. This platform supports a variety of functional assays and provides a reproducible framework for studying microglial reactivity in Alzheimer’s disease and related disorders. Its modularity allows for future applications involving Aβ or tau stimulation, complement modulation, or therapeutic testing, making it a versatile tool for dissecting microglial contributions to synaptic pathology.

## Declarations

### Ethics approval

All animal procedures were approved by the local animal welfare committees and conducted in accordance with the guidelines of the German Research Council (DFG) and the European Directive 2010/63/Eu and according to the French law (APAFIS authorization number 59-350-009), as appropriate.

### Consent for publication

Not applicable.

### Availability of data and materials

The transcriptomic datasets generated during this study are available from the authors upon request.

### Authors’ contributions

**DK** and **JCL** conceived and supervised the study, supported data interpretation, secured funding, and critically reviewed the manuscript. **DSW** coordinated the project, performed most experiments, analyzed data, and wrote the manuscript. **AMA** provided essential technical support, including development of the co-culture system, preparation and maintenance of primary cultures, and animal colony management. **CL, FE, VB**, and **JC** contributed technical support at different stages of the study. **DB, DH**, and **MD** performed library preparation and RNA sequencing. **LI** analyzed transcriptomic data. **KB** performed microfabrication. **MP, KPK**, and **TM** generated the Hexb-tdTomato mouse line. All authors read and approved the final manuscript.

## Acknowledgements

We thank the BioImaging Center Lille (BiCeL) and La PLateforme d’Expérimentation et de Haute Technologie Animale (PLEHTA), both part of Plateformes Lilloises en Biologie et Santé (PLBS) - UAR 2014 - US 41), for microscopy maintenance and animal housing, respectively.

